# FcɛR1γ-based Activating Chimeric Antigen Receptors enhanced Natural Killer cell function against HLA-E^+^ cells

**DOI:** 10.1101/2025.06.16.659984

**Authors:** Lewis L. Lanier, John R. Greenland, Frank Momburg, Daniel R. Calabrese, Avishai Shemesh

## Abstract

FcɛR1γ is a transmembrane adaptor protein regulating NKp30, NKp46, and CD16 expression and function in human NK cells. HLA-E expression by cancer cells suppresses NK cell function through interaction with NKG2A. Here, we engineered human NK cells to express a modified FcɛR1γ with a 4-1BB co-stimulatory motif. FcɛR1γ^41BB^ expression led to upregulation of NKp30, NKp46, and CD16 expression and enhanced NKp30 and CD16-mediated target cell lysis. Against cells expressing HLA-E, FcɛR1γ^41BB^ improved NK cell lytic function relative to FcɛR1γ. We then generated an CD19^ScFv^-CD8^hinge^-FcεR1γ^41BB(TM-IC)^ CAR (FCRG10). Engineering human NK cells to express FCRG10 led to upregulation of NKp30 and CD16, and improved NK cell-mediated cytotoxicity against cells co-expressing B7H6, CD19, and HLA-E relative to NK cells expressing the conventional CD19CAR^41BB-CD3ζ^ with a CD8 transmembrane domain. Our results indicate that modification of FcɛR1γ enhanced NK cell lytic function mediated by several activating receptors and improved CAR NK cell response in conditions favoring NK cell suppression by HLA-E. Our work may contribute to NK cell-based cancer immunotherapy mediated by co-recognition of stress-induced ligands, antibody-coated cells, and tumor-associated antigens.

## Introduction

The expression of chimeric antigen receptors (CAR) in immune effector cells is a promising therapeutic strategy for cancer treatment^1,2^. CARs allow the recognition of tumor-associated antigens that lead to effector cell activation and target cell killing through immunoreceptor tyrosine-based activation motif (ITAM) phosphorylation^3^. First-generation CARs were designed by fusion of a single-chain fragment variable (ScFv) chain with the CD3ζ or FcɛR1γ transmembrane (TM) adapter proteins, which contain three or one ITAMs, respectively, in their intracellular (IC) domain^4^. T cells expressing the first-generation CAR-CD3ζ^TM-IC^ displayed a superior signaling efficacy and effector functions relative to CAR-FcɛR1γ^TM-IC4^. However, the CAR-CD3ζ^TM-IC^ forms complexes with the endogenous TCR/CD3 complex, which reduces the CAR’s stability due to competition with the native TCR complex and leads to higher tonic signaling^5,6^. Thus, second and third generation CARs integrated other hinge and transmembrane domains, such as CD8α, to improve the CAR’s stability and to reduce non-specific T cell activation through interactions with the endogenous TCR/CD3 complex^6,7^. In 2017, Tisagenlecleucel, the CD19^ScFv^CD8^hinge-TM^41BB^IC^CD3ζ^IC^ CAR (CD19-CAR^41BB-CD3ζ^), targeting human CD19, was the first CAR T cell therapy to be approved by the Food and Drug Administration (FDA) for the treatment of B-cell acute lymphoblastic leukemia (ALL) or diffuse large B-cell lymphoma (DLBCL)^2^.

Natural killer (NK) cells are innate lymphocytes with an innate ability to kill cancer cells directly and are used for cancer immunotherapy^8–10^. CD19-CAR NK cells show promising clinical efficacy in phase 1 and 2 trials in patients with CD19-positive lymphoid tumors^11^. Unlike CAR T cell therapy, CAR NK cell therapy does not lead to cytokine release syndrome, neurotoxicity, graft-versus-host disease, or increased IL-6 levels, and it can be used in allogeneic treatments^12^. Several sources of human NK cells, such as NK cell lines (e.g., NK92)^13^, primary cord blood or peripheral blood NK cells, or iPSC differentiated NK cells, provide potential “off-the-shelf” therapy^12^. There are several ongoing NK cell-based clinical trials for treating hematopoietic and non-hematopoietic malignancies^12,14^. Still, the conventional CAR constructs designed for T cells may not be optimal for NK cells^2^.

NK cells’ innate ability to eliminate cancer cells is orchestrated by an array of germline-encoded activating receptors that recognize stress-induced self-ligands or antibody-coated cells^8^. An array of inhibitory receptors recognizing HLA class I molecules suppress NK cell activation and leads to tolerance^15^. One of the main inhibitory receptors expressed by NK cells is NKG2A, encoded by the *KLRC1* gene^16,17^. Human NKG2A recognizes several leader peptide-HLA-E complexes^18^. The upregulation of activating receptor or stress-induced self-ligands overcomes inhibition mediated by HLA class I recognition^15^. Activating receptors upregulation and activation occur through transmembrane interactions with several transmembrane adaptor proteins^19^. Unlike T cells, which mainly do not express FcɛR1γ^20^, both FcɛR1γ and CD3ζ are expressed by primary NK cells. Homo- or heterodimers of CD3ζ or FcɛR1γ^21^ regulate the surface expression and activation signaling of the activating receptors NKp30, NKp46, or CD16, through transmembrane interaction, while DAP10 regulates the expression of NKG2D, and DAP12 regulates NKG2C and NKp44 expression^19^. Recently, several groups reported that loss of FcɛR1γ protein expression in primary human NK cells leads to loss of NKp30 and NKp46 surface protein expression and impairs NK cell function against cancer cells^22–25^. Still, FcɛR1γ loss leads to a mild decrease in CD16 protein expression and increases CD16 antibody-dependent function through interaction with CD3ζ homodimers^25,26^. This suggests that human FcɛR1γ has a higher binding affinity than CD3ζ to NKp30, NKp46, or CD16 transmembrane domains. Therefore, we hypothesized that modification of FcɛR1γ intracellular domain with a co-stimulatory motif would increase NKp30, NKp46, and CD16 expression in human CD3ζ^+^ NK cells, and lead to enhanced activating receptor-dependent NK cell innate and adaptive-like lytic function.

## Results

### FcɛR1γ^41BB^ expression leads to NKp30, NKp46, or CD16 upregulation

To test our hypothesis, we first expressed the native form of FcɛR1γ or FcɛR1γ^41BB^ in the CD3ζ^+^ FcɛR1γ^negative^ CD16^negative^ human NK cell line, NK92 (Figure 1A, 1B, S1)^10,27^ and examined if FcɛR1γ^41BB^ would upregulate NKp30, NKp46, and CD16 expression, similar to FcεR1γ ability. Expression of FcɛR1γ or FcɛR1γ^41BB^ in NK cells did not change the expression levels of CD3ζ (Figure 1C). NKp30 or NKp46 surface protein expression increased by FcɛR1γ or FcɛR1γ^41BB^ (Figure 1D), indicating that FcɛR1γ expression leads to upregulation of NKp30 and NKp46. In line, *FCER1G* CRISPR knockout led to a relative decreased of NKp30 or NKp46 expression (Figure 1E) in comparison to the parental cells or fluorescent minus one (FMO) staining control. This confirmed the upregulation of NKp30 and NKp46 is FcɛR1γ-dependent and indicates that basal NKp30 or NKp46 expression observed in parental cells is due to CD3ζ. Surface protein expression of the inhibitory receptor NKG2A, the activating receptors NKG2D or NKp44, or the activation markers 4-1BB^28^ or TRAIL^29^ was not affected by FcɛR1γ or FcɛR1γ^41BB^ expression (Figure 1F). To test the effect of FcɛR1γ^41BB^ expression on CD16 expression, we engineered parental NK92 cells to express human CD16 (Figure 2A). A CD107a assay using mouse IgG1 antibody-coated beads against CD16 or other activating receptors confirmed that CD16 is functional, indicating it can associate with endogenous CD3ζ in NK92 cells (Figure 2B), without affecting the function of NKp30, NKp46, or NKG2D (Figure 2B). Expression of FcɛR1γ or FcɛR1γ^41BB^ in CD16^+^ NK92 cells led to upregulation of CD16 surface expression relative to parental NK cells, as was observed for NKp30, used as a positive control (Figure 2C). Thus, our results indicate that FcεR1γ^41BB^ leads to upregulation of NKp30, NKp46, or CD16, similar to FcεR1γ ability.

**Figure 1:**
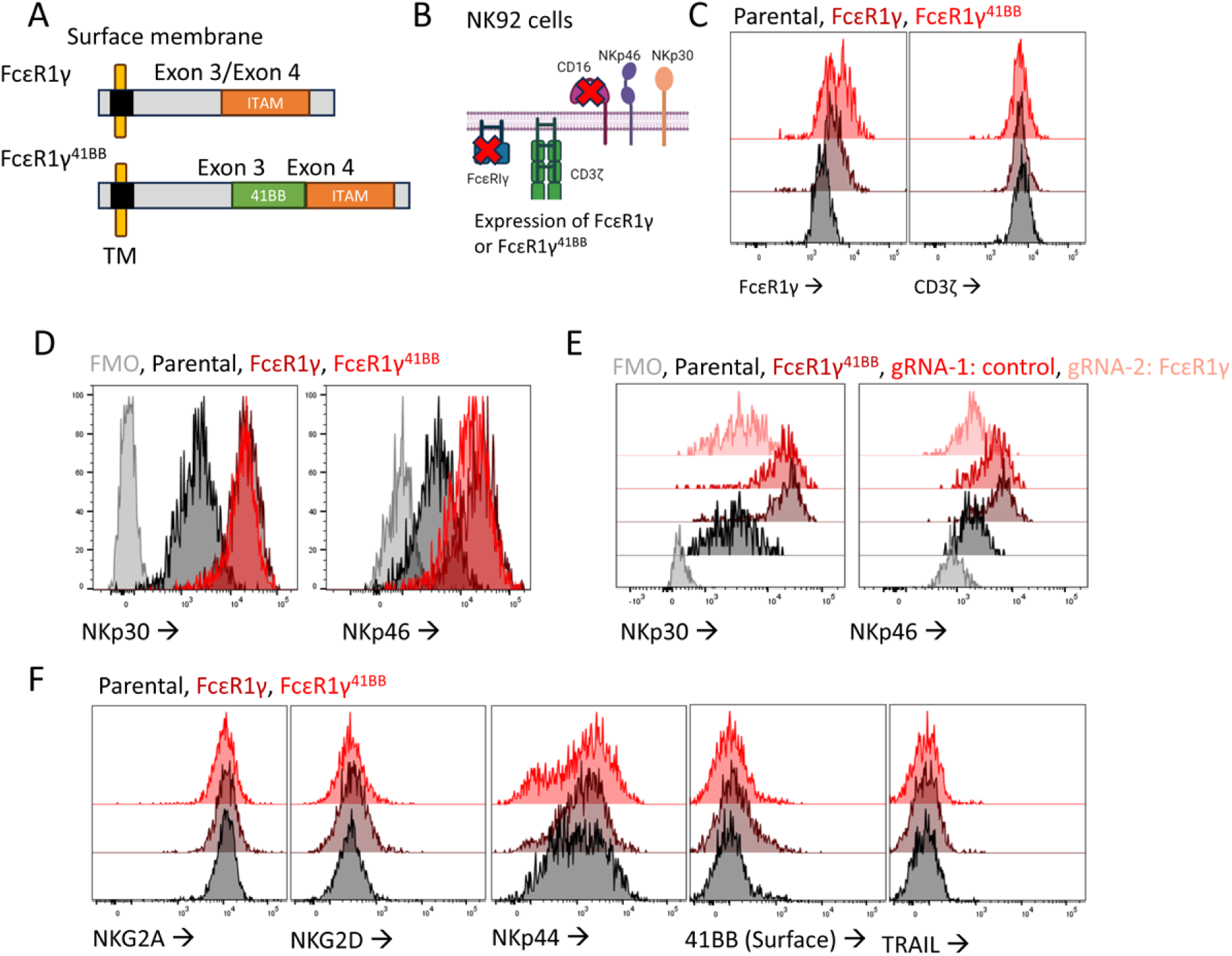
FcεR1γ^41BB^ increases NK cell NKp30 and NKp46 expression. A) Schematic representation of FcɛR1γ^41BB^ construct design relative to FcɛR1γ native form. B) Schematic representation of relevant activating receptors and adaptors expressed in the natural killer cell line, NK92. C) Flow-cytometry histograms of FcɛR1γ or CD3ζ intracellular protein expression in the FcεR1γ or FcεR1γ^41BB^ transduced NK92 cells relative to parental cells (color-coded). D) Flow-cytometry histograms of NKp30 or NKp46 protein surface expression in the FcεR1γ or FcεR1γ^41BB^ transduced NK92 cells relative to parental cells or fluorescent minus one (FMO) control. (color-coded). E) Flow-cytometry histograms of NKp30 or NKp46 protein surface expression in the transduced FcɛR1γ^41BB^ NK cells following *FCER1G* gene CRISPR-knockout relative to parental cells or a fluorescent minus one (FMO) control. (color-coded). F) Flow-cytometry histograms for the surface protein expression of the inhibitory receptor NKG2A, the activating receptor NKG2D or NKp44, or the activation markers 41BB or TRAIL in NK92 cells (color-coded).

**Figure 2:**
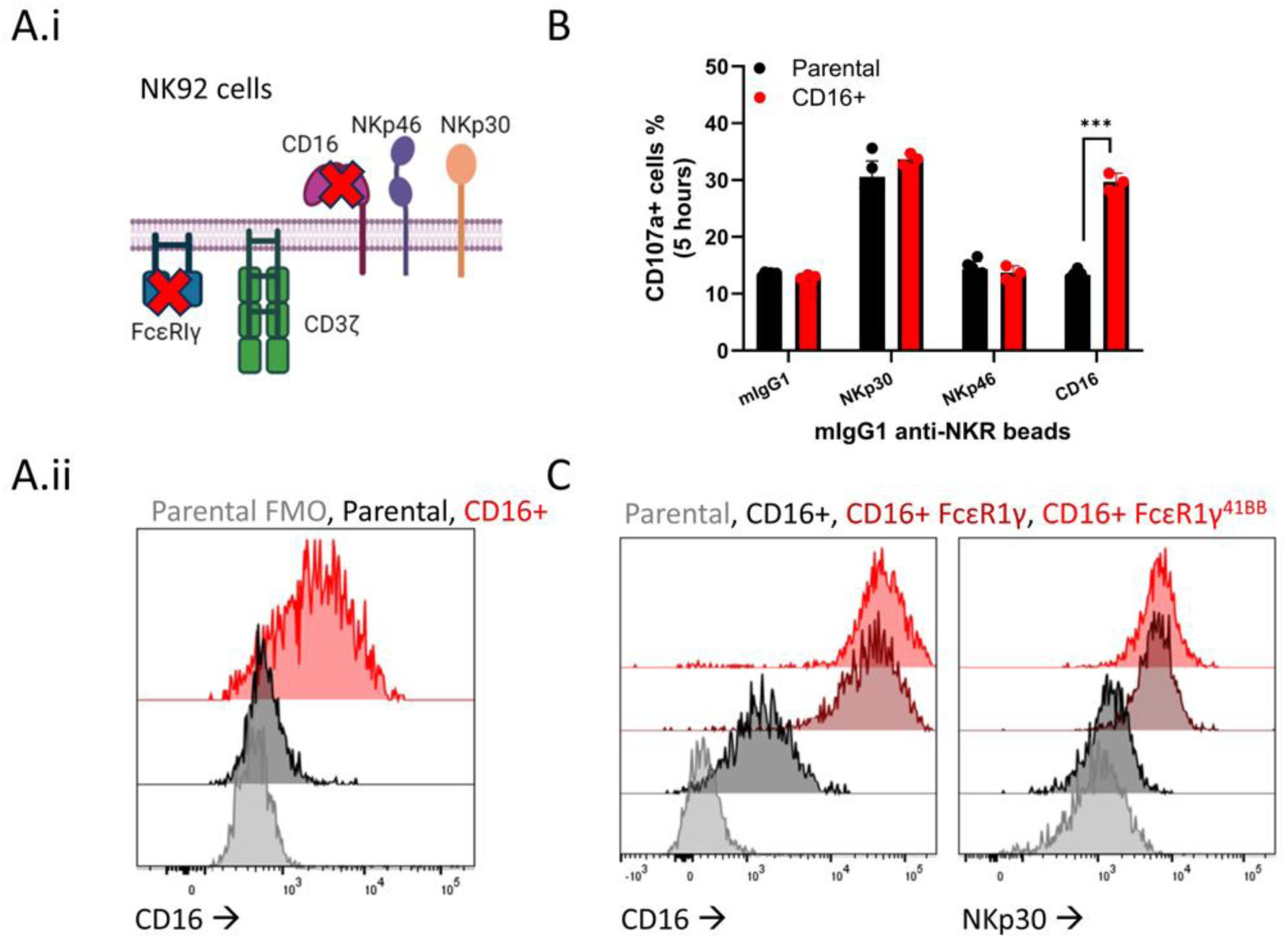
FcεR1γ^41BB^ increases NK cell CD16 expression. A) Schematic representation of relevant activating receptors and adaptors expressed in the natural killer cell line, NK92 (i). Flow-cytometry histograms of CD16 surface expression in CD16 transduced or parental NK cells and relative to fluorescent minus one FMO control (color-coded). B) CD107a assay: CD16^+^ NK cells or parental NK cells were stimulated for 5 hours with anti-NKp30, anti-NKp46, anti-CD16, or anti-NKG2D mIgG1 coated beads relative to mIgG1 control. Mean +/- SEM, unpaired t-test, two tails. NK receptors (NKR). C) Flow-cytometry histograms of CD16 or NKp30 in NK cells (color-coded).

### FcɛR1γ^41BB^ increases NKp30-dependent NK cell function

We hypothesized that FcεR1γ^41BB^, relative to FcεR1γ, would mediate an increase in activating receptor-dependent NK cell lytic function against target cells. Thus, we performed a 24-hour killing assay at low effector to target (E:T) ratios against chronic myelogenous leukemia (CML) cell line, K562, known to express the NKp30 ligand, B7H6, with low to negative expression of HLA-E^30–32^. Activating receptor-human Fc fusion protein binding assay confirmed the binding of NKp30-hFc and NKG2D-hFc to K562 cells (Figure S2A). Additionally, B7H6 staining of K562 cells, relative to B7H6^+^ Ba/F3 cells^33^ used as a positive control, confirmed the expression of B7H6 on K562 cells (Figure S2B). FcɛR1γ^+^ or FcɛR1γ^41BB+^ NK cells mediated higher K562 cell lysis. (Figure 3A.i) FcɛR1γ^41BB+^ NK cells displayed a 20 percent increase in target cell lysis of B7H6^+^NKG2D-ligand^+^HLA-E^-^ K562 cells at lower effector to target ratios relative to FcɛR1γ^+^ NK cells, which may suggest the contribution of NKG2D co-stimulation during NKp30-mediated activation^34^. B7H6 knockout in K562 cells using CRISPR-Cas9 resulted in minimal target cell lysis with no differences between parental, FcɛR1γ^+^, or FcɛR1γ^41BB+^ NK92 cells (Figure 3A.ii), thus confirming the increase in K562 killing was NKp30-dependent and the contribution of other weaker mechanisms leading to K562 cell lysis by NK92 cells^34^. To further test NKp30-dependent lysis, we performed a 5-hour killing assay using B7H6^+^ or parental Ba/F3 cells. FcɛR1γ^+^ or FcɛR1γ^41BB+^ NK cells did not mediate non-specific target cell lysis of parental Ba/F3 cells (Figure 2B.i). Against B7H6^+^ Ba/F3 cells, FcɛR1γ^41BB+^ NK cells displayed an increased target cell lysis relative to FcɛR1γ^+^ or parental NK cells (Figure 2B.ii). These data suggest FcɛR1γ^41BB^ increases NKp30-dependent NK cell cytotoxicity. Additionally, we observed that FcɛR1γ^+^ NK92 cells showed no difference in target cell lysis relative to parental cells (Figure 2B.ii). This suggested that FcɛR1γ may block NKp30 interaction with CD3ζ homodimers or time-dependent target cell lysis through NKp30 in the presence of FcɛR1γ due to lower ITAM numbers. These results confirmed that FcεR1γ^41BB^ increases NKp30-dependent NK cell lytic function against B7H6^+^ target cells.

**Figure 3:**
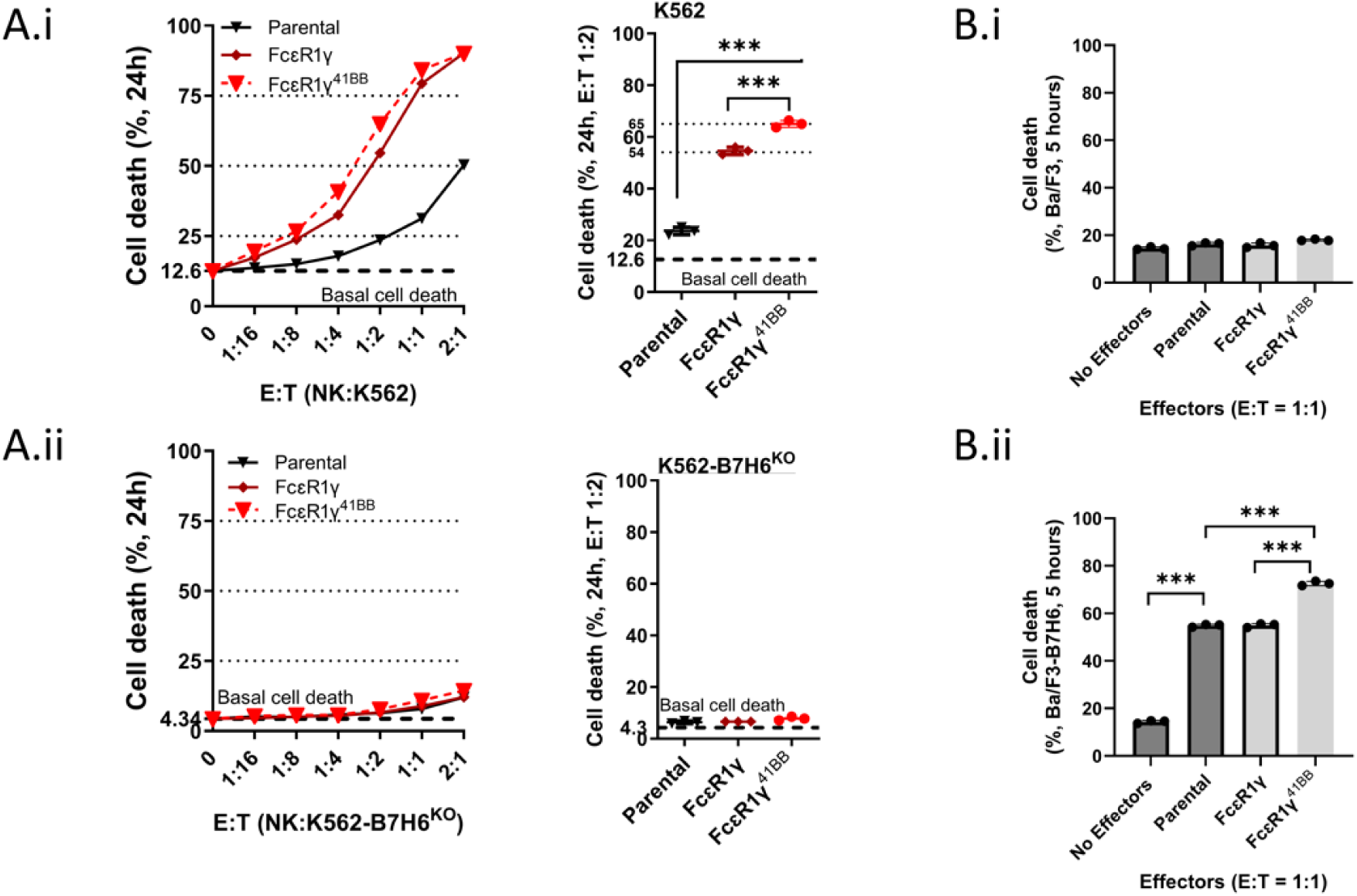
FcεR1γ^41BB^ increases NKp30 function. A) Killing assay: K562 (A.i) or K562 B7H6-knockout (A.ii) cells were co-incubated for 24 hours with by parental, FcɛR1γ, or FcεRIγ^41BB^ NK cells at the indicated E:T ratios. K562 cell death was measured by propidium iodide (PI) staining. Mean+/ SEM, unpaired t-test, one-tailed, **p < 0.001, ***p < 0.001. The dashed line showing Basal K562 cell death with no effectors. B) Killing assay: Ba/F3 (i) or B7H6^+^ Ba/F3 cells (ii) cells were co-incubated for 5 hours with by parental, FcɛR1γ, or FcεR1γ^41BB^ NK cells at the indicated E:T ratio. Ba/F3 cell death was measured by propidium iodide (PI) staining. Mean+/ SEM, unpaired t-test, one-tailed, **p < 0.001, ***p < 0.001.

We examined NKp46-dependent degranulation by CD107a assay using anti-human NKp46 mouse IgG1-coated beads. Anti-NKp30-coated beads were used as positive control (Figure S2C). NKp30 stimulation increased NKp30-dependent activation and degranulation of FcɛR1γ^41BB+^ NK92 cells. NKp46 stimulation of FcɛR1γ^41BB+^ NK cells did not increase CD107a expression (Figure S2D). In contrast, anti-NKp46 coated beads stimulation of IL-2 cultured primary NK cells lead to a higher degranulation (Figure S2D), while NKp46 expression in FcɛR1γ or FcɛR1γ^41BB+^ NK92 cells was similar to NKp46 expression in IL-2 cultured primary NK cells (Figure S2E).

### FcɛR1γ^41BB^ increases CD16-dependent NK cell function

To test if CD16-mediated function was increased by FcɛR1γ^41BB^ relative to FcɛR1γ, we performed a CD107a assay. FcɛR1γ^+^ or FcɛR1γ^41BB+^ CD16^+^ NK cells were stimulated with anti-human CD16 mouse IgG1-coated beads. FcɛR1γ^+^ or FcɛR1γ^41BB+^ CD16^-^ NK cells were used as a negative control. Following CD16 cross-linking, FcɛR1γ^41BB+^ CD16^+^ NK cells displayed a significant increase in CD107a surface expression relative to FcεR1γ^+^ or parental CD16^+^ cells (Figure 4A.i). Stimulation of CD16^-^ NK cells did not increase CD107a surface expression, thus confirming CD16-dependent activation and degranulation (Figure 4A.ii). To test CD16 function during human antibody Fc recognition, we engineered Ba/F3 cells to express human CD20 (Figure 4B) and tested for antibody-dependent cellular cytotoxicity (ADCC) using Rituximab (Figure 4C). FcɛR1γ^41BB+^ CD16^+^ NK92 cells displayed a significant increase in ADCC relative to FcεR1γ^+^ or parental CD16^+^ NK92 cells (Figure 4D). FcɛR1γ^+^ cells showed no difference relative to parental cells, suggesting FcɛR1γ may block CD16 interaction with CD3ζ homodimers or time-dependent lysis of target cells by CD16 in the presence of FcɛR1γ. We concluded that FcεR1γ^41BB^ increases the CD16-mediated function of CD3ζ^+^ NK cells.

**Figure 4:**
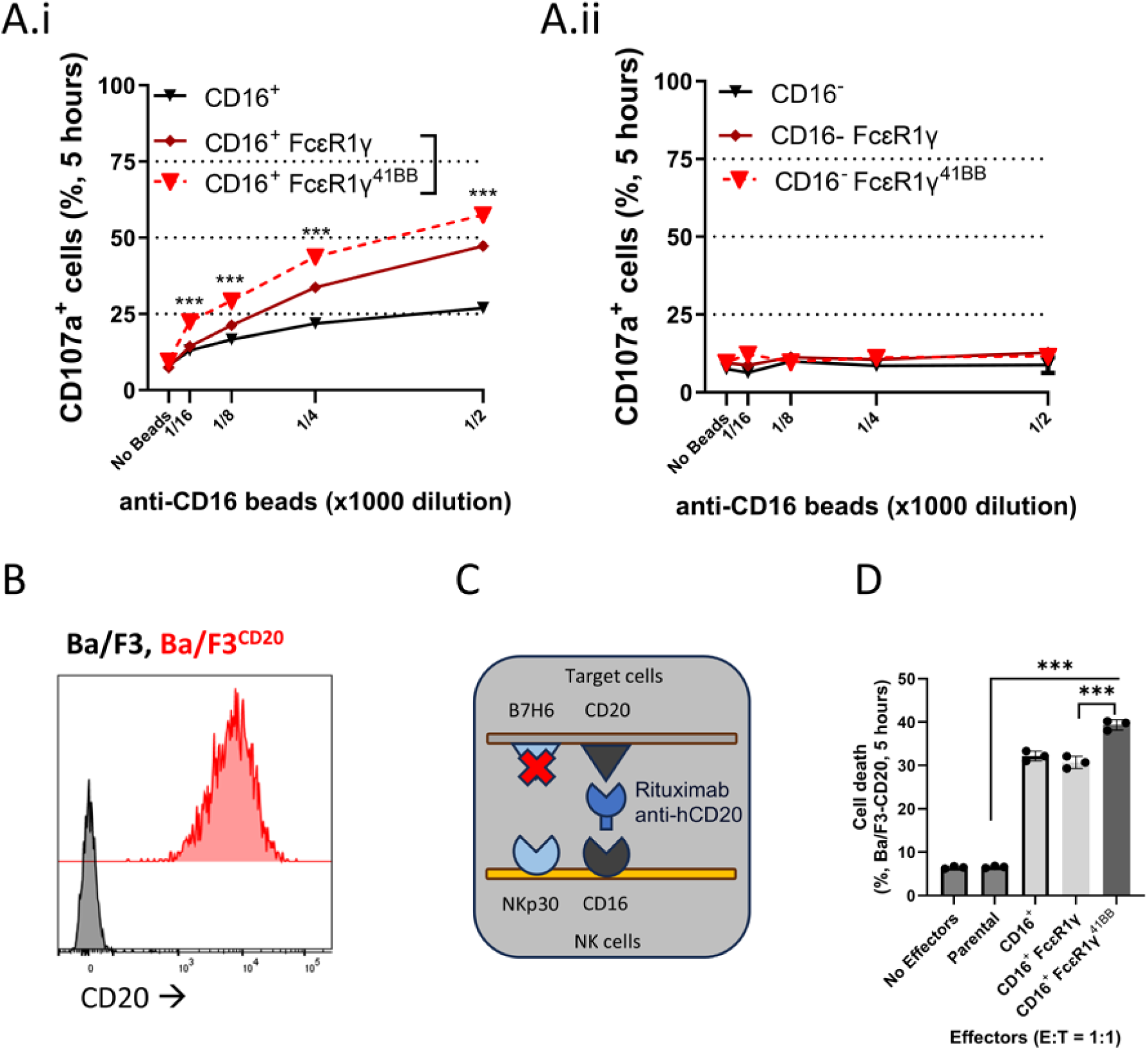
FcεR1γ^41BB^ increases NK cell CD16 function. A) CD107a assay: CD16^+^ (black), CD16^+^ FcɛR1γ (dark red), or CD16^+^ FcɛR1γ ^41BB^ (red) NK cells were stimulated for 5 hours with anti-CD16 beads (A.i). CD16^-^ (black), CD16^-^ FcɛR1γ (dark red), or CD16^-^ FcɛR1γ^41BB^ (red) NK cells were stimulated for 5 hours with anti-CD16 beads (A.ii). Mean+/- SEM, unpaired t-test, two-tailed, ***p < 0.001. t-test was calculated between CD16^+^ FcɛR1γ vs. CD16^+^ FcɛR1γ ^41BB^ NK cells. B) Flow-cytometry histograms of human CD20 expression in Ba/F3 cells(color-coded). C) Schematic representation of killing assay by ADCC using Rituximab (anti-human CD20) in our Ba/F3 model. D) Killing assay: NK cells were co-cultured with Ba/F3-CD20 cells for 5 hours at an (E:T ratio of 1:1. Ba/F3 cell death was assessed with propidium iodide (PI) staining. Mean+/ SEM, unpaired t-test, one-tailed, ***p < 0.001.

### FcɛR1γ^41BB^ increases NK cell function against HLA-E^+^ cells

Targeting NKG2A-HLA-E interaction is essential for cancer immunotherapy^35^. We hypothesized that FcɛR1γ^41BB^ would improve NK cell lytic function against cells co-expressing B7H6 and HLA-E. Therefore, we co-expressed peptide-tethered HLA-E molecules with different leader peptides (HLA-C or HLA-G) in B7H6^+^ Ba/F3 cells (Figure S3A). In line with HLA-E-presented HLA-G leader peptide being the strongest ligand for NKG2A^36,37^, HLA-E^HLA-G^ peptide complexes elicited the most significant decrease in NK cell function in the presence of B7H6 (Figure S3B), which was NKG2A-dependent as confirmed by NKG2A knockout (Figure S3C). Therefore, we tested the NKp30-dependent NK cell lysis function of FcɛR1γ^41BB+^ NK cells against B7H6^+^HLA-E^HLA-G^ Ba/F3 cells at an E:T ratio of 1:1 for 5 hours (Figure 5A, 5B). FcɛR1γ^41BB+^ NK cells displayed significantly higher target cell-mediated lysis relative to FcɛR1γ-expressing cells or parental FcɛR1γ^-^ cells. This indicates that FcɛR1γ^41BB^ expression promotes a superior NK cell lytic function in the presence of HLA-E, independently of NKp30 expression levels. Additionally, FcɛR1γ^+^ cells displayed a higher target cell lysis relative to parental FcɛR1γ^-^ cells. This suggested that in the presence of HLA-E, higher NKp30 surface expression levels play a positive role in overcoming NKG2A-mediated inhibition during a short co-incubation.

**Figure 5:**
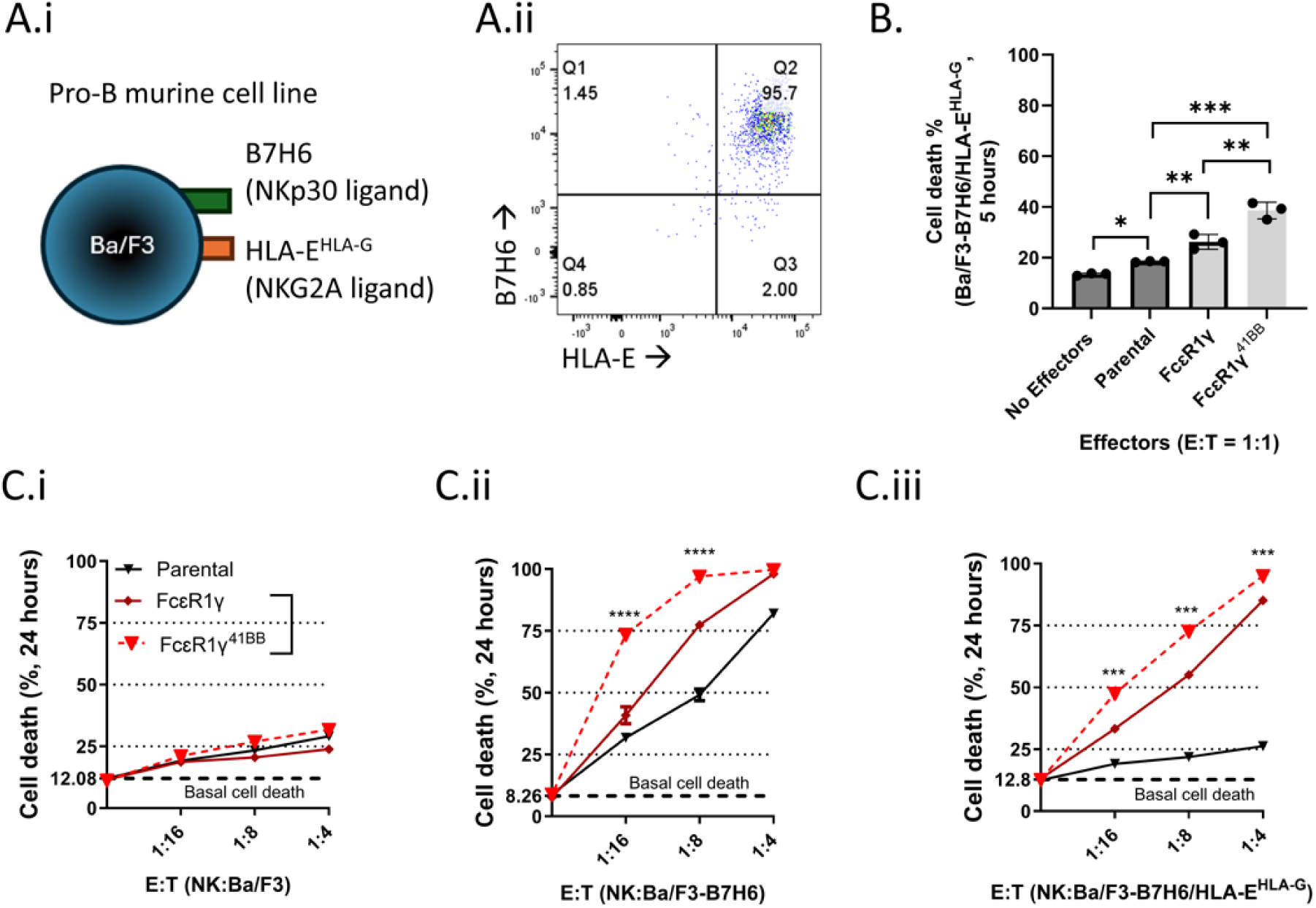
FcεR1γ^41BB^ increases NK cell NKp30 function against HLA-E^+^ cells. A) Schematic representation of the pro-B murine cell line, Ba/F3, engineered to express B7H6 or human HLA-E (A.i). Flow-cytometry dot-plot panel of the expression of B7H6 vs. human HLA-E (A.ii). B) Killing assay: B7H6^+^HLA-E^+^ Ba/F3 cells (right) cells were co-incubated for 5 hours with parental, FcɛR1γ, or FcɛR1γ^41BB^ NK cells at the indicated E:T ratio. Ba/F3 cell death was measured by propidium iodide (PI) staining. Mean+/ SEM, unpaired t-test, two-tailed, *p < 0.05, **p < 0.001, ***p < 0.001. C) Killing assay: parental (B.i), B7H6^+^ (B.ii), or B7H6^+^HLA-E^+^ Ba/F3 cells were co-incubated for 24 hours with by parental, FcɛR1γ, or FcɛR1γ^41BB^ NK cells at the indicated E:T ratios. Ba/F3 cell death was measured by propidium iodide (PI) staining. Mean+/ SEM. The dashed line showing basal K562 cell death with no effectors. Mean+/ SEM, unpaired t-test, two-tailed, ***p < 0.001, ****p < 0.0001.

To test our observations in setting more favorable to the HLA-E-mediated inhibition, we performed a 24-hour killing assay at lower E:T ratios against B7H6^+^HLA-E^+^ Ba/F3 cells and used B7H6^+^ or parental Ba/F3 cells as controls (Figure 5C). FcɛR1γ^41BB+^ NK cells displayed a significantly higher target cell lysis of B7H6^+^ or B7H6^+^HLA-E^+^ Ba/F3 cells relative to FcɛR1γ^+^ NK cells. This result confirmed that FcɛR1γ^41BB^ expression can enhance NK cell NKp30-dependent lytic function against B7H6^+^HLA-E^+^ cells. Additionally, FcɛR1γ^+^ NK cells displayed a significantly higher target cell lysis than parental cells against B7H6^+^ or B7H6^+^HLA-E^+^ Ba/F3 cells. This indicated that in the presence of HLA-E, higher NKp30 expression levels due to FcɛR1γ expression 1) can overcome NKG2A-mediated NK cell tolerance more efficiently, 2) significantly enhance target cell lysis in a time-dependent manner relative to CD3ζ homodimers in FcɛR1γ^-^cells.

### CD19^ScFv^-CD8^hinge^ fused to FcεR1γ^41BB^ increases NKp30 and CD16 expression

Chimeric antigen receptors (CARs) are used in cancer immunotherapy. Thus, co-expression of a CAR and FcεR1γ^41BB^ may enhance NK cell function against cancer cells co-expressing tumor-associated antigens and B7H6 or ligand recognition by an antibody. However, we hypothesized that fusion of FcεR1γ^41BB^ with an extracellular ScFv domain of a CAR will provide the CAR the ability to increase FcεR1γ-dependent activating receptors expression and function (Figure S1D). For that purpose, we fused the CD19^ScFv^-CD8^hinge^ extracellular domain of a Myc-tag-CD19^ScFv^-CD8^hinge-TM^-41BB^IC^CD3ζ^IC^ CAR (CD19CAR^41BB-CD3ζ^) with FcεR1γ^41BB^ transmembrane and intracellular domains (from exon 2 which contains the first cysteine for adaptor’s dimerization^19^, FCRG10, Figure 6A). FCRG10 expression led to upregulation of NKp30 and CD16, as was detected in FcεR1γ^41BB^-expressing cells (Figure 6B). Relative to FCRG10, CD19CAR^41BB-CD3ζ^ transduced CD16^+^ NK cells did not display upregulation of NKp30 or CD16 expression (Figure 6B.i). The increase in NKp30 or CD16 expressions in FCRG10^+^ NK cells corresponded to the expression of Myc-tag, located in the N-terminus of the CAR (Figure 6B.ii). This shows that fusion of FcεR1γ^41BB^ with an extracellular CD19^ScFv^-CD8^hinge^ domain can upregulate NKp30 and CD16 expression.

**Figure 6:**
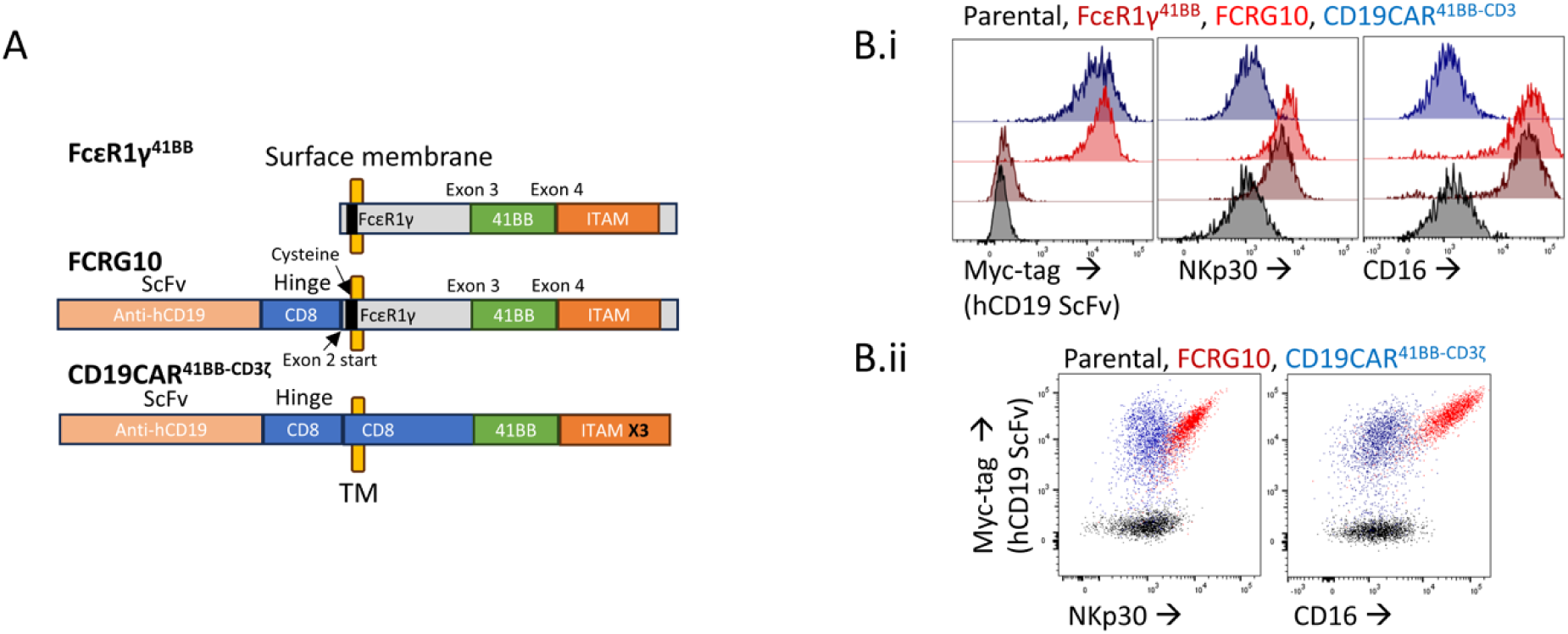
CD19^ScFv^-CD8^hinge^ fused to FcεR1γ^41BB^ increases NKp30 and CD16 expression. A) Schematic representation of CD19^ScFv^CD8^hinge^ fused to FcεR1γ^41BB^ (FCRG10; ITAMs X1) design relative to FcɛR1γ^41BB^ or CD19CAR^41BB-CD3ζ^ (ITAMs X3) design. A Myc-tag was integrated at the N-terminus of FCRG10 or CD19CAR^41BB-CD3ζ^ for detection of surface expression. B) Flow-cytometry histograms of Myc-tag, NKp30 or CD16 in CD16^+^ NK cells (Parental (black), FcɛR1γ^41BB^ (dark red), FCRG10 (red), or CD19CAR^41BB-CD3ζ^ (blue) (B.i). Flow-cytometry dot-plots of NKp30 or CD16 surface expression relative to Myc-tag expression (B.ii).

### CD19^ScFv^-CD8^hinge^-FcεR1γ^41BB^ CAR improves NK cell function against HLA-E^+^ target cells

To test the function of FCRG10, we engineered Ba/F3, Ba/F3-B7H6^+^, or Ba/F3-B7H6^+^HLA-E^+^ cells to express human CD19 (Figure 7A) and did a killing assay side-by-side (Figure 7B). FCRG10^+^ or CD19CAR^41BB-CD3ζ^ ^+^ NK cells show no non-specific target cell lysis of parental Ba/F3 cells relative to other controls (Figure 6B.i). Against CD19^+^ Ba/F3 cells, FCRG10^+^ or CD19CAR^+^ NK cells displayed high target cell lysis, indicating that FCRG10 is functional (Figure 6B.ii). Still, FCRG10^+^ NK cells displayed a higher target cell lysis, possibly due to the transmembrane dimerization of FcεR1γ with CD3ζ. Against CD19^+^B7H6^+^ Ba/F3 cells, FCRG10^+^ NK cells displayed increased target cell lysis relative to FcεR1γ^41BB+^ NK cells (Figure 6B.iii). This suggested that FCRG10 mediates improved recognition of two activating ligands and increased NK cell-mediated target cell lysis. FCGR10^+^ NK cells showed no difference from CD19CAR^+^ NK cells (Figure 6B.iii). This suggests the recognition of two activating ligands, by a CAR and NKp30, and in the absence of HLA-E, functionally compensates for the differences in NKp30 expression levels mediated by FcεR1γ expression. Against CD19^+^B7H6^+^HLA-E^+^ Ba/F3 cells, FCRG10^+^ NK cells displayed a significant increase in target cell lysis relative to CD19CAR^+^ NK cells (Figure 6B.iv). This indicates that in the presence of HLA-E and dual activating ligand recognition (B7H6 and CD19), the increased NKp30 expression and function mediated by the FCRG10 enhance NK cell lytic function relative to a CD19CAR^41BB-CD3ζ^.

**Figure 7:**
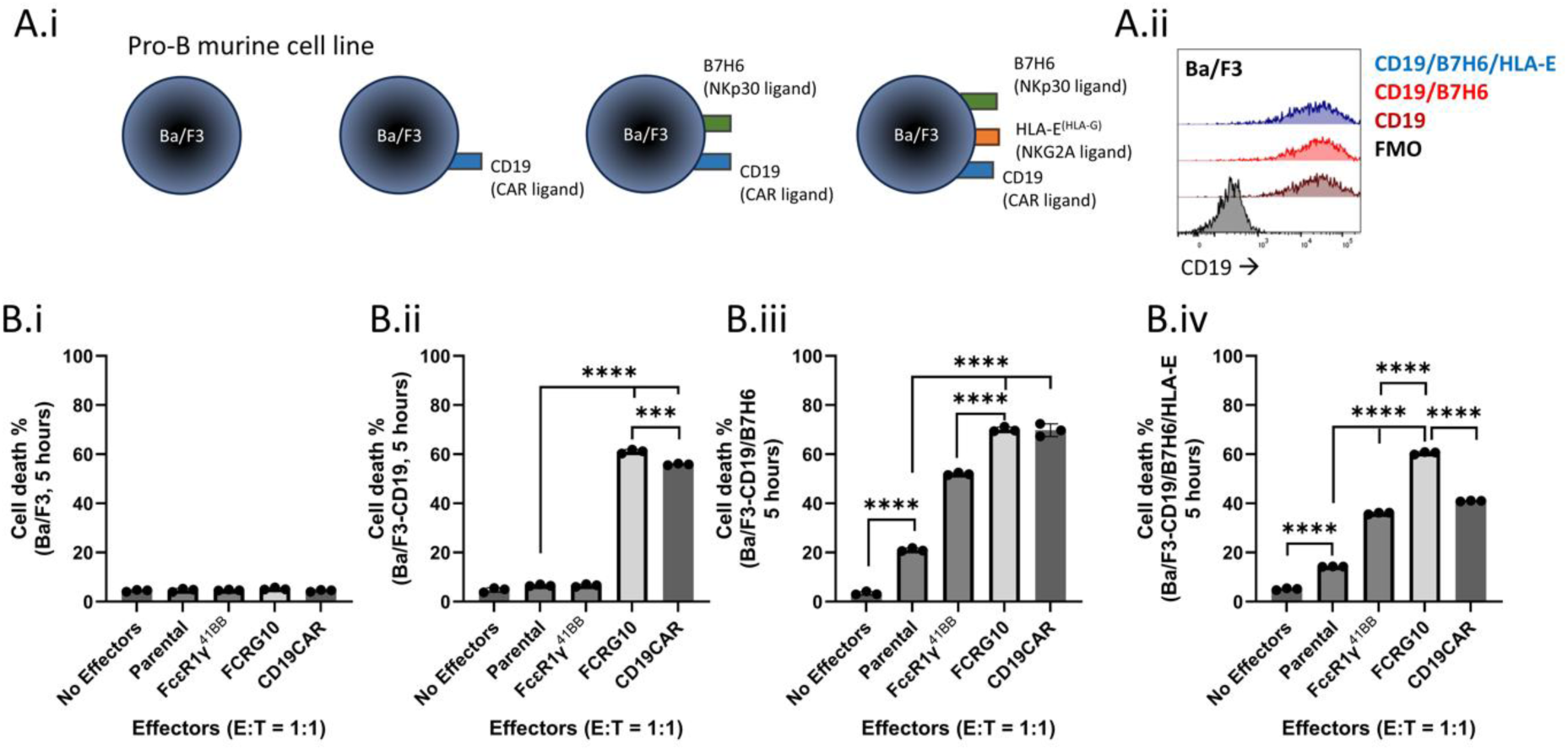
CD19^ScFv^-CD8^hinge^-FcεR1γ^41BB^ fusion improves NK cell function against HLA-E^+^ target cells. A) A.i) Schematic representation of Ba/F3 cells engineered to co-express human CD19, B7H6 and HLA-E. A.ii) Flow-cytometry histograms of human CD19 surface expression in Ba/F3 cells. B) Killing assay: parental (B.i), human CD19^+^ (B.ii), human CD19^+^B7H6^+^ (B.iii, or human CD19^+^B7H6^+^HLA-E^+^ (B.iv) Ba/F3 cells were co-incubated for 5 hours with by parental, FcɛR1γ^41BB^, FCRG10, or CD19CAR NK92 cells at a E:T ratio of 1:1. Ba/F3 cell death was measured by propidium iodide (PI) staining. Mean+/ SEM, unpaired t-test, two-tails, ***p < 0.001, ****p < 0.0001.

## Discussion

In this study, we hypothesized that expression of a modified FcεR1γ would lead to upregulation of NKp30, NKp46, and CD16 through transmembrane interaction and increase NK cell lytic function mediated by these activating receptors. We have shown that expression of a modified FcɛR1γ adaptor with a 4-1BB co-stimulatory motif increases human NK cell-mediated lysis and degranulation mediated by the activating receptors NKp30 and CD16. Interestingly, our results showed that NKp46 stimulation through anti-NKp46 coated beads did not result in increased NK92 cell degranulation, even though NKp46 was upregulated by FcεR1γ expression. This might suggest an additional mechanism of regulation for NKp46 function^38^ or that NKp46 function in NK92 cells is impaired, and should be further investigated with a NKp46-ligand expressing cells. Thus, our study shows that FcεR1γ expression leads to upregulation of NKp30, NKp46, and CD16. Furthermore, modification of FcεR1γ with 4-1BB increases the function of several activating receptors. Therefore, FcεR1γ^41BB^ increases the innate ability of NK cells to eliminate cancer cells expressing stress-induced self-ligands and mimics the increased CD16 function observed in primary human FcεR1γ^-^ NK cells^25^. This is important as the ability of NK cells to eliminate cancer cells is regulated by the integrated signals from several activating receptors, especially when the ligand expression is low^15^. Additionally, the use of ADCC against several cancer-associated antigens (e.g., human CD20 (Rituximab)^39^, human HER2 (Trastuzumab)^40^, human EGFR (Cetuximab)^41^, and others), or with bispecific antibodies (e.g., AFM13 targeting CD30 and CD16)^42^, or NK cell engagers (NKCEs, targeting NKp46 and CD16, and a tumor antigen)^43^, has shown promising results in pre-clinical and clinical trials. Thus, engineering NK cells with a modified FcεR1γ may additionally complement these therapeutic approaches for cancer immunotherapy.

We have shown that modification of FcεR1γ improves NK cell target cell lysis of HLA-E^+^ cells in the presence of an activating ligand by increasing the activating receptor function and expression. This is important as NK cells provide allogeneic “off the shelf” therapeutic benefit for cancer immunotherapy and express several inhibitory receptors recognizing HLA class-I molecules (e.g., KIR2DL1-DL5, KIR3DL1-DL3, or NKG2A). Due to the allogeneic nature of NK cell therapy, KIR-HLA-A/B/C mismatches rates are high, and therefore, targeting NKG2A-HLA-E interaction is essential for cancer immunotherapy. Targeting the NKG2A-HLA-E axis by NKG2A blocking antibody (e.g., Monalizumab)^35^ or by CRISPR knockout of *KLRC1*^16^ improves NK cell anti-cancer function in pre-clinical and clinical trials. Still, others have shown that NKG2A knockout diminished human NK cell expansion^44^, and in the absence of MHC class-I recognition, NK cells become hyporeactive^45^. Thus, modification of FcεR1γ can be a new approach to improves NK cell function and overcomes tolerance mediated by the NKG2A-HLA-E axis through activating receptor upregulation and enhanced function during ligand recognition and may maintain NK cell reactivity and expansion capacity. Additionally, the activating receptor NKG2C is triggered by the same pHLA-E complexes as NKG2A^46^. Thus, engineering NKG2C^+^ NK cells with a modified FcεR1γ may improve their lytic function through co-recognition of HLA-E and self-induced ligands, such as B7H6, or antibody coated cells.

We generated an anti-CD19 CAR with a modified FcɛR1γ^TM-IC^ (FCRG10). We have shown that relative to the conventional anti-CD19 CAR design, the FCRG10 is functional and enhances NK cell function against HLA-E^+^ target cells co-expressing NKp30 and CD19-CAR ligands. Hence, tailored CAR design for NK cells can improve the cells’ anti-cancer function. Additionally, higher NKp30 protein expression is associated with a positive outcome of multiple myeloma (MM)^47^ or acute myeloid leukemia (AML)^48^ patients. As CD19-CAR NK cells showed promising results in clinical trails^11^, using FCRG10 or FcɛR1γ^41BB^ fused to other ScFv chains recognizing BCMA or CD33 may improve the innate ability of NK cells to eliminate cancer cells in human leukemia patients. As the construct size of FcɛR1γ^41BB^ is only 387 base-pairs, an additional approach is to co-express a CAR and a modified FcɛR1γ.

Overall, our study shows that FcɛR1γ modification or CAR-FcɛR1γ construct design are tailored for human NK cells and can enhance their function against cancer cells co-expressing self-induced ligands, antibody-dependent recognition of tumor-associated ligands, and CAR-dependent tumor-associated antigens.

## Methods

### Cloning and lentiviral particles preparation

Human FcɛR1γ cDNA (Sequence ID: NM_004106.2), FcɛR1γ^41BB^, Myc-tag-CD19^ScFv^CD8^hinge^FcεR1γ^41BB(TM-IC)^ (FCRG10), human CD16 cDNA (Sequence ID: NM_000569.8), human CD19 cDNA (Sequence ID: NM_001178098.2), human CD20 cDNA (Sequence ID: NM_152866.3), Myc-tag-CD19^ScFv^-CD8^hinge-TM^-41BB^IC^-CD3ζ^IC^ chimeric antigen receptor (conventional CD19CAR^41BB-CD3ζ^, generously provided by Dr. K. T. Roybal, University of California, San Francsico, USA), or pHLA-G- or pHLA-C-β2M-HLA-E constructs were cloned into the lentivirus vectors pHR containing the SFFV promotor using an In-Fusion® HD Cloning Kit (TAKARA). FcɛR1γ contained exons 1-5 of the NM_004106.2 cDNA. FcɛR1γ^41BB^ contained exons 1-5 and was designed by inserting a human 41BB co stimulatory motif (i.e., KRGRKKLLYIFKQPFMRPVQTTQEEDGCSCRFPEEEEGGCEL, similar to 41BB motif found in the Myc-tag-CD19^ScFv^-CD8^hinge-TM^-41BB^IC^-CD3ζ^IC^ CAR) between exon 3 and exon 4 of *FCER1G* cDNA sequence. FcɛR1γ^41BB^ overall size; 129 amino acids, 387 base-pairs. FCRG10 contained nucleic acids 1-954 of CD19^ScFv^-CD8 ^hinge^-41BB-CD3ζ chimeric antigen receptor (CAR) ectodomain (e.g., CD8 signal peptide, Myc-tag, CD19^ScFv^, and CD8^hinge^) fused to transmembrane and intracellular domains of FcɛR1γ^41BB^ at amino acid # 17 (e.g., from *FCER1G* exon 2, which contains the first cysteine and transmembrane domain of FcɛR1γ). DNA fragments were obtained from Integrated DNA Technologies (e.g., IDT). Membrane-bound HLA-E single-chain trimer constructs HA1L-HLA-C(V3-L11)GC-, or HA1L-HLA-G(V3-L11)GC-*GSL12*-hβ_2_m-*GSL21*-AviTag-HLA-E*01:03-ED[Y84C]-A*02:01-TMD/ICD _T2A_IgκL-BirA_KDEL_ were generously provided by Dr. F. Momburg, German Cancer Research Center (DKFZ), Heidelberg, Germany). HLA-C leader peptide sequence: VMAPRTLIL, HLA-G leader peptide sequence: VMAPRTLFL.

Lentivirus preparation was done by using the pMD2.G and pCMV dr8.91 packaging vectors and transfection of the Lenti-X™ 293T cell Line (TAKARA, cat. 632180) cultured in complete DMEM plus 10% FCS. Lentivirus was concentrated using a Lenti-X™ concentrator (TAKARA, cat. 631232) and resuspended in 1 ml RPMI-1640 + 10% fetal calf serum (FCS) with protamine sulfate (1 μg/ml). Aliquots were kept at −20°C.

### Cell culture and transduction

NK92 cells (CRL-2407) were obtained from “The American Type Culture Collection” (ATCC). NK92 cell culture media: Minimum Essential Medium Eagle (Sigma Aldrich, cat. M0200-500ML) + 20% horse serum (Sigma Aldrich, cat. H1138) + human IL-2 (25 ng/ml, Peprotech, cat 200-02-1MG). Mouse pro-B cell line Ba/F3 pre-engineered to express mouse IL-3 were generated by Dr. Lewis L. Lanier. Ba/F3 or K562 culture media: complete RPMI-1640 + 10% heat-inactivated FCS. For lentiviral transductions, 50,000 cells were mixed with concentrated lentiviral particles in 96-well round-bottom plate (200 µl of culture media) and were centrifuged at 1000 RCF for 1 hour at room temperature. After centrifugation, cells were incubated at 37° C, 5% CO2) for 24 hours. Cells were then transferred to T25 flask for expansion. NK cells were then sorted to match the expression of NKp30, NKp46, or CD16, between FcɛR1γ, FcɛR1γ^41BB^, or FCRG10 transduced cells. Ba/F3 cells were sorted to match the expression of indicated ligands.

### Flow cytometry and sorting

The following antibodies or fusion proteins were used for the detection of cell surface receptors, or intracellular proteins, and NK cell receptor binding. Anti-FcεR1γ (Millipore, cat. FCABS440F), anti-CD3ζ (BioLegend, cat. 644103), anti-NKp30 (BioLegend, cat. 325208, cat. 325210), anti-NKp46 (BioLegend, cat. 331908, cat. 331936), anti-NKG2A (Miltenyi Biotech, act. 130-113-563), anti-NKG2D (R&D, cat. FAB139P), anti-NKp44 (BioLegend, cat. 352112), anti-41BB (BioLegend, cat. 309809), anti-TRAIL/CD253 (BioLegend, cat. 308209), anti-B7H6 (R&D, cat. FAB7144P, cat. FAB7144A), anti-HLA-E (BioLegend, cat. 342605. eBioscience, cat, 12-9953-71), anti-CD16 (BioLegend, cat. 302012, cat. 302015, cat. 302038), anti-CD19 (BioLegend, cat. 302208, cat. 302242), anti-Myc-tag (Cell Signaling, cat 3739S), hNKG2D-hFc (R&D, cat. 1299-NK), hNKp30-hFc (R&D, cat. 1849-NK), hNKp44-hFc (R&D, cat. 2249-NK), hNKp46-hFc (R&D, cat. 1850-NK), hFc (R&D, cat. 110-HG), and anti-human IgG (Jackson, cat. 109-117-008, cat. 109-236-170). Staining and wash buffer: PBS + 2% FCS. For surface antigen detection, antibodies were incubated with the cells for 30 minutes at 4°C. For receptor-human Fc chimera surface staining, proteins were incubated with the cells for 60 minutes at 4° C, washed off and then cells were incubated with anti-human IgG for 30 minutes at 4° C. For intracellular antigen detection, cells were incubated for 20 min at 4°C with 100 µl/well Cytofix/Cytoperm buffer (51-2090KZ; Becton Dickinson). Following incubation, cells were washed twice using Perm Wash buffer (cat. 421002; BioLegend) diluted 1:10 in PBS and then the antibodies were incubated with the cells for 60 minutes at room temperature. Protein expression, or co-expression was detected by flow cytometry LSR-II; Becton Dickinson Immunocytometry Systems. Engineered FcɛR1γ, FcɛR1γ^41BB^, FCRG10, or CD19CAR NK92 cells were sorted by the increased expression or co-expression of NKp30, NKp46, CD16, or Myc-tag. Ba/F3 cells were sorted by the expression or co-expression of B7H6, HLA-E, CD20, CD19 using FACSAria: Becton Dickinson Immunocytometry Systems. Dead cells were excluded by using Zombie NIR™ Fixable Viability Kit (BioLegend, cat. 423106) diluted 1:500 in PBS + 2% FCS.

### CRISPR-Cas9 knockout of *FCER1G, KLRC1* (NKG2A), or *NCR3LG1* (B7H6)

FcεR1γ^41BB^-positive or parental NK92 cells or K562 cells were edited using the non-viral CRISPR-Cas9 system. crRNA was resuspended in IDT duplex buffer at a final concentration of 160 µM. crRNA and tracrRNA, obtained from IDT, were mixed at a 1:1 volume ratio and were incubated at 37 °C for 30 min to form gRNA solution. gRNA solution and Cas9-NLS (Berkeley QB3 MacroLab) were mixed at a 1:1 volume ratio and were incubated at 37 °C for 15–30 min. Cells were then resuspended in 18 µl Lonza P3 buffer + Cas9/gRNA solution and were then transferred to the Lonza 96-well electroporation shuttle. Electroporation codes: NK92 cells; CM137, K562 cells; FF120. *FCER1G*: crRNA-1 control; GCCCAAGATGATTCCAGCAG, *FCER1G*: crRNA-2 specific: CAGCTCTGCTATATCC TGGA. *NCR3LG1*: crRNA; AAGTAGAGATGATGGCAGGG, or ATGGTGACAT TGTCATTCAG. *KLRC1:* crRNA: AGGAGTAATCTACTCAGACC

### Antibody-conjugated beads and cell stimulation

Antibody-conjugated beads were prepared according to the company’s protocol (Invitrogen Dynabeads Antibody Coupling Kit, cat. 14311D) at 10 µg antibody per 1 mg beads. Following conjugation, beads were resuspended in sterile PBS at an antibody concentration of 0.1 µg/µl. Antibody conjugation was evaluated by flow cytometry with APC-conjugated anti-mouse or rat IgG. BioLegend: anti-CD16 (cat. 302002, IgG1k), anti-NKp30 (cat. 325204, IgG1k), (cat. 325102, IgG1k), anti-NKp46 (cat. 331904, IgG1k), anti-NKG2D (cat. 320802, IgG1k), and mouse IgG1k isotype-matched control (cat. 400102). Antibody-conjugated beads were kept at 4°C until further use. The assays were performed in 96 round-bottom well plates. Antibody-coated beads were diluted at 1:1,000 or as indicated in NK92 cell culture media and used by adding 50 µl/well. NK92 cells (5 × 10^4^ cells/well) were added at 150 µl/well in NK92 cell culture media. Alexa Fluor 647 anti-human CD107a antibody (BioLegend, cat. 328620) was added at the beginning of the assay at a final dilution of 0.1 µl/well. NK92 cells were incubated at 37°C with 5% CO_2_ for the duration of the assay. Cells were analyzed for the detection of CD107a by flow cytometry (LSR-II; Becton Dickinson Immunocytometry Systems). Dead cells were excluded by using propidium iodide (1 mg/ml, 1:500). NK92 cells incubated without beads were used for CD107a background control and the increase in CD107a signal was calculated relative to each cell line.

### NK cell lysis assay and ADCC

NK92 cells were labeled with cell trace violet (CTV) according to the company’s protocol (Invitrogen, cat. C34557;). Ba/F3 or K562 cells (target cells) were labeled with CFSE (Invitrogen, cat. C34554). The assays were done only when the viability of the NK92 cells was above 90% as measured by Trypan Blue and performed in 96 well round-bottom plates. NK92 cells were resuspended in NK92 cell culture at a concentration of 1 × 10^5^ cells/100 µl / well, and were diluted 1:2 1:4 or as indicated in NK92 cell culture media. Target cells were resuspended in complete RPMI-1640 + 10% heat-inactivated FCS, at a concentration of 5 × 10^4^ cells/ 100 µl / well. Cell co-cultures were incubated at 37°C with 5% CO2 for the assay duration. For ADCC, anti-human Rituximab (anti-CD20, Selleckchem, cat. A2009) was added to the co-culture at a final concentration of 5 µg/ml. At the end of the assay, cells were stained using propidium iodide (1 mg/ml, 1:500) for the detection of dead cells and percentage of dead cells was calculated as PI^+^CFSE^+^ relative to PI^-^CFSE^+^ cells while CTV^+^ cells were excluded. Samples were analyzed by flow cytometry (LSR-II; Becton Dickinson Immunocytometry Systems).

### Graphics and statistical analysis

Graphs were generated using GraphPad Prism 10 or FlowJo_V10.8.1 Statistical analysis is indicated in figure legends and was calculated using GraphPad Prism 10 or Excel (Microsoft 365). *, P < 0.05; **, < P < 0.01; ***, P < 0.001, **** P < 0.0001.

## Acknowledgments

Studies were supported by the Parker Institute for Cancer Immunotherapy. PICI Award Number: C-03171. Institutional Reference Number: A125828. We thank Dr. Kole T. Roybal for providing us with the CD19CAR^41BB-CD3ζ^ plasmid.

Author contributions: Conceptualization (A. Shemesh); Data curation (A. Shemesh); Formal analysis (A. Shemesh); Funding acquisition (A. Shemesh, L. L. Lanier, J. R. Greenland, D.R. Calabrese); Investigation (A. Shemesh); Methodology (A. Shemesh); Project administration (A. Shemesh); Resources (A. Shemesh, F. Momburg, L.L. Lanier); Supervision and Validation (A. Shemesh); Visualization (A. Shemesh); original draft (A. Shemesh); review & editing (A. Shemesh, L.L. Lanier, F. Momburg, J. R. Greenland).

## Supplementary Figures

**Supplementary Figure 1:**
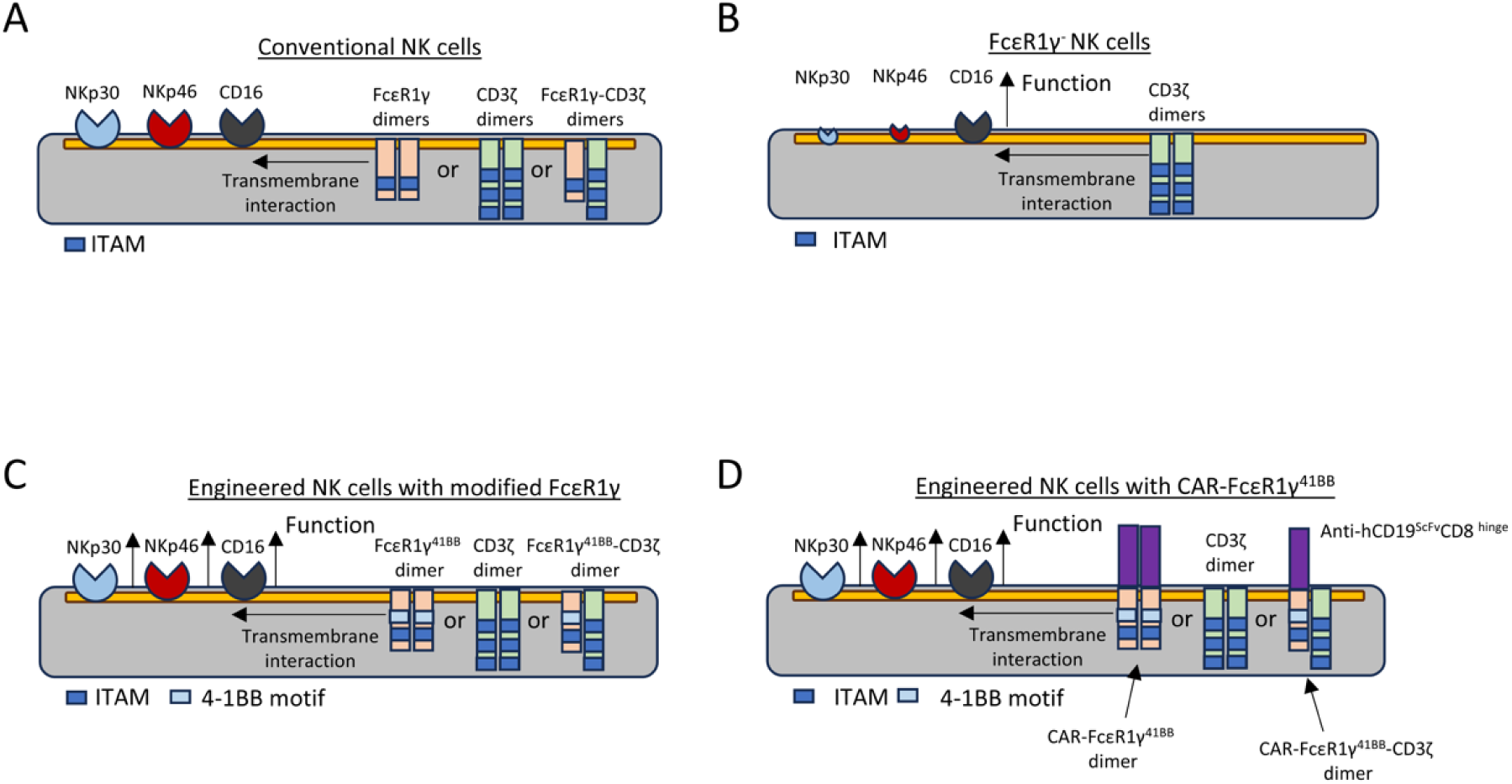
Schematic representation of study aims. A) Human conventional NK cells express the adaptor proteins FcεR1γ or CD3ζ in a homo- or heterodimer forms. These dimers regulate the expression and function of NKp30, NKp46, and CD16 through transmembrane interaction with the activating receptor. B) FcεR1γ knockout leads to loss of NKp30 and NKp46 expression, decrease CD16 expression. CD16-mediated NK cell function increases relative to conventional NK cells. C) Engineered NK cells with a modified FcεR1γ. NK cells were engineered to express a modified FcεR1γ (FcεR1γ^41BB^) to test if this approach would 1) increase NKp30, NKp46, and CD16 expression and 2) increase NKp30- and NKp46-, and CD16-mediated NK cell function relative to in FcεR1γ^-^CD3ζ^+^ or in FcεR1γ^+^CD3ζ^+^ NK cells. D) Engineering NK cells with a CAR-FcεR1γ^41BB^. NK cells were engineered to express an anti-hCD19^ScFv^CD8^hinge^FcεR1γ^41BB(TM-IC)^ to test if this approach would increase activating receptor expression in a CAR-dependent manner and improve NK cell lytic function in the presence of several activating ligands.

**Supplementary Figure 2:**
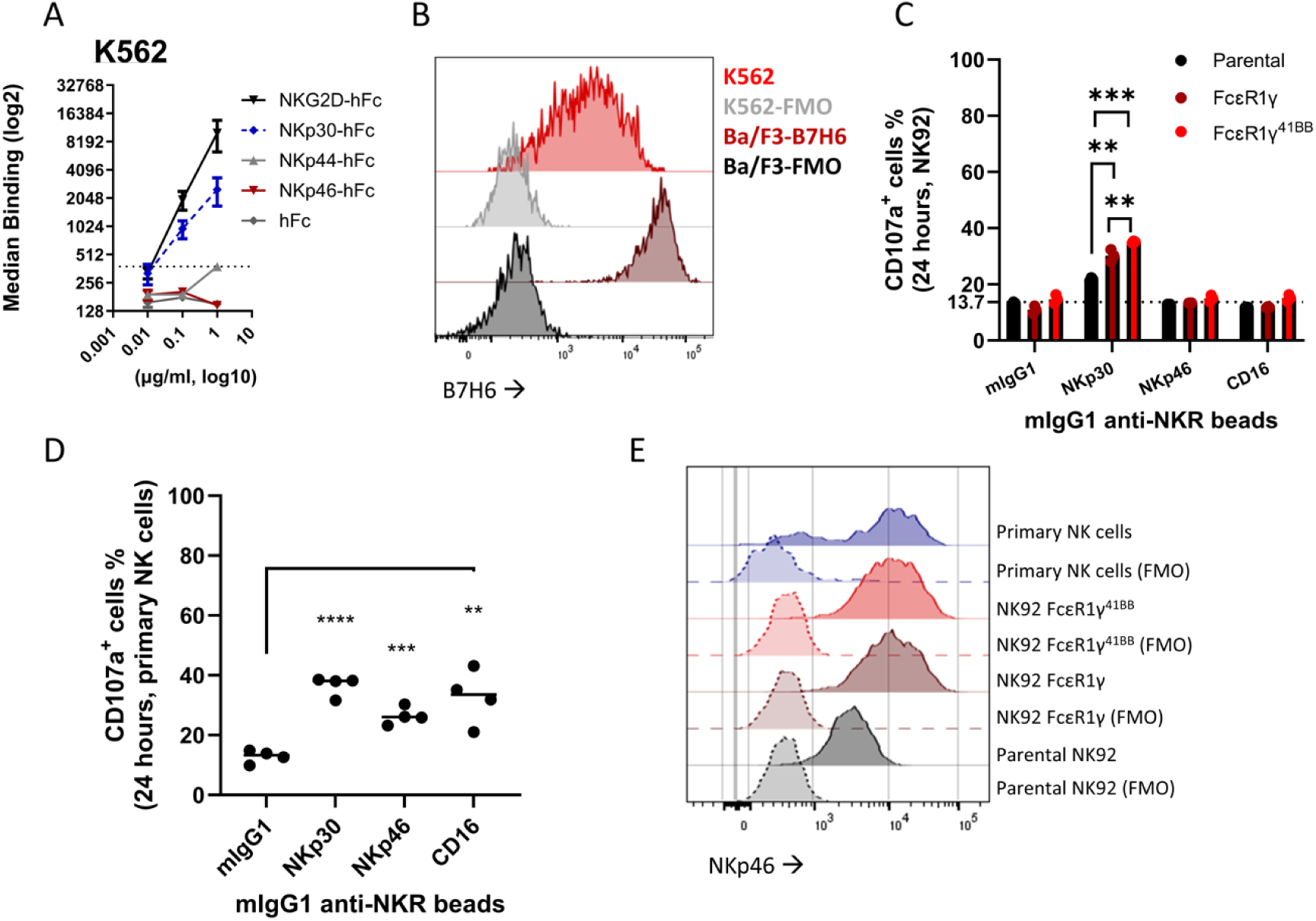
B7H6 expression on K562 cells and NKp46-mediated function. A) Median binding values of NK cell receptor-human Fc fusions to the human AML cell line, K562. B) Flow-cytometry histograms of B7H6 surface expression in Ba/F3 cells or K562 cells relative to florescent minus one (FMO) controls. C) CD107a assay: NK cells were stimulated for 24 hours with anti-NKp30, anti-NKp46, or anti-CD16 coated beads relative to mIgG1 control in the presence of IL-2. Mean +/- SEM, unpaired t-test, two tails. ** p < 0.01, *** p < 0.001. NK receptors (NKR). D) CD107a assay: IL-2 cultured primary NK cells were stimulated for 24 hours with anti-NKp30, anti-NKp46, or anti-CD16 coated beads relative to mIgG1 control in the presence of IL-2. Mean +/- S.D, unpaired t-test, two tails. ** p < 0.01, *** p < 0.001, **** p < 0.0001. NK receptors (NKR). E) Flow cytometry histograms with grids showing NKp46 expression on NK92 cells (parental, FcεR1γ, or FcεR1γ^41BB+^) relative to IL-2 cultured primary NK cells and FMO controls.

**Supplementary Figure 3:**
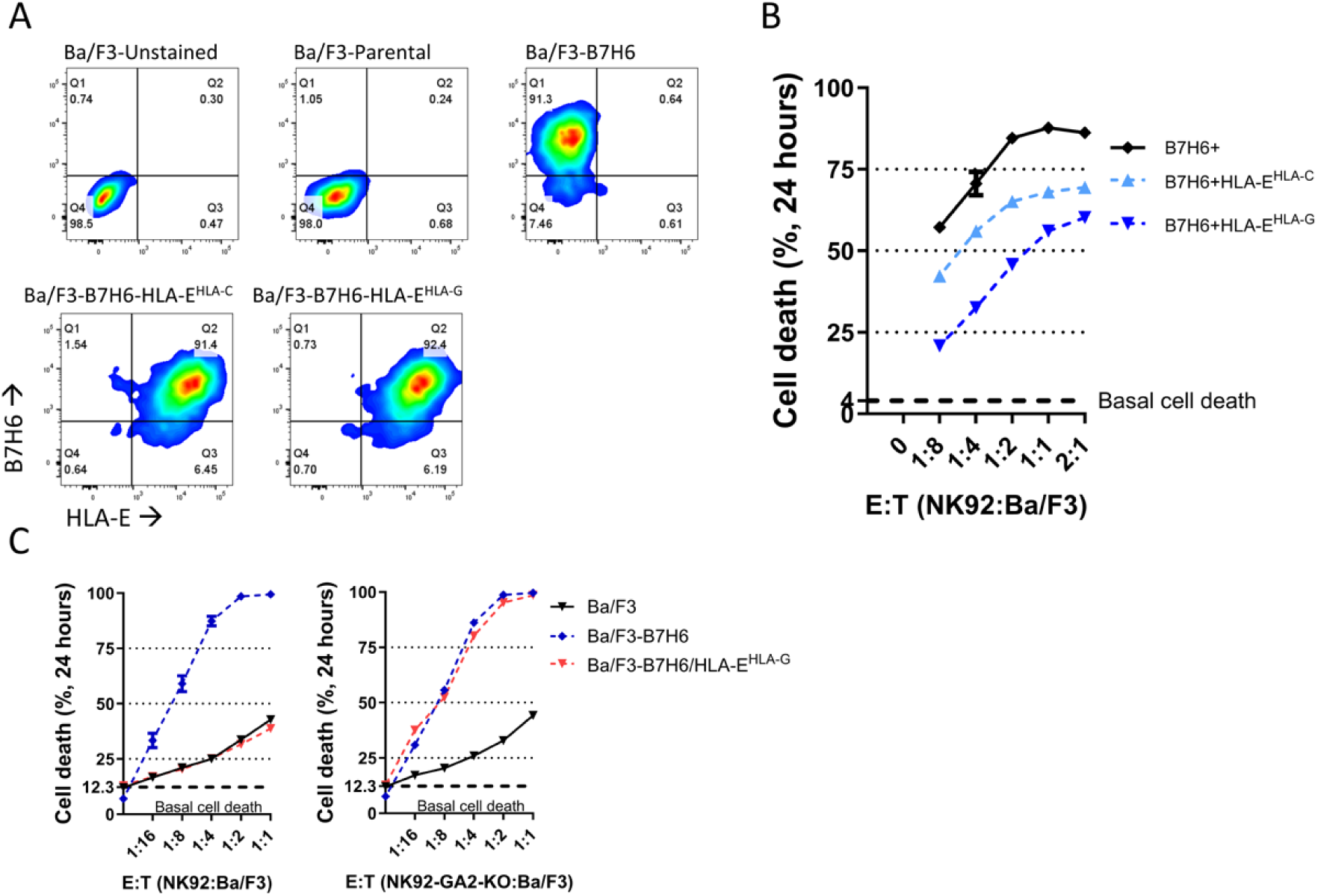
HLA-E-mediated NKG2A-dependent NK cell suppression. A) Flow-cytometry plots of Ba/F3 cells, engineered to express B7H6 and HLA-E with HLA-C, or HLA-G leader peptide. B) Inhibition of NK cell NKp30-dependent killing of Ba/F3 cells by HLA-E expression during B7H6 co-expression. C) Influence of NKG2A knockout (G2A-KO) on HLA-E^HLA-G^ mediated suppression of NK cell lysis of Ba/F3 cells during B7H6 co-expression.

## Notes

### Competing Interest Statement

The authors have declared no competing interest.

## References

1. Sterner, R. C. & Sterner, R. M. CAR-T cell therapy: current limitations and potential strategies. Blood Cancer J. 11, 1–11 (2021).

2. Ruppel, K. E., Fricke, S., Köhl, U. & Schmiedel, D. Taking Lessons from CAR-T Cells and Going Beyond: Tailoring Design and Signaling for CAR-NK Cells in Cancer Therapy. Front. Immunol. 13, (2022).

3. Wu, L., Wei, Q., Brzostek, J. & Gascoigne, N. R. J. Signaling from T cell receptors (TCRs) and chimeric antigen receptors (CARs) on T cells. Cell Mol Immunol 17, 600–612 (2020).

4. Haynes, N. M. et al. Redirecting Mouse CTL Against Colon Carcinoma: Superior Signaling Efficacy of Single-Chain Variable Domain Chimeras Containing TCR-ζ vs FcεRI-γ1. The Journal of Immunology 166, 182–187 (2001).

5. Bridgeman, J. S. et al. The optimal antigen response of chimeric antigen receptors harboring the CD3zeta transmembrane domain is dependent upon incorporation of the receptor into the endogenous TCR/CD3 complex. J Immunol 184, 6938–6949 (2010).

6. Dwivedi, A., Karulkar, A., Ghosh, S., Rafiq, A. & Purwar, R. Lymphocytes in Cellular Therapy: Functional Regulation of CAR T Cells. Front. Immunol. 9, (2019).

7. Fujiwara, K. et al. Hinge and Transmembrane Domains of Chimeric Antigen Receptor Regulate Receptor Expression and Signaling Threshold. Cells 9, 1182 (2020).

8. Cerwenka, A. & Lanier, L. L. Natural killer cell memory in infection, inflammation and cancer. Nat Rev Immunol 16, 112–123 (2016).

9. Shimasaki, N., Jain, A. & Campana, D. NK cells for cancer immunotherapy. Nat Rev Drug Discov 19, 200–218 (2020).

10. Zhang, B. et al. Chimeric antigen receptor-based natural killer cell immunotherapy in cancer: from bench to bedside. Cell Death Dis 15, 1–18 (2024).

11. Liu, E. et al. Use of CAR-Transduced Natural Killer Cells in CD19-Positive Lymphoid Tumors. New England Journal of Medicine 382, 545–553 (2020).

12. Wang, W. et al. Breakthrough of solid tumor treatment: CAR-NK immunotherapy. Cell Death Discov. 10, 1–16 (2024).

13. Klingemann, H. The NK-92 cell line—30 years later: its impact on natural killer cell research and treatment of cancer. Cytotherapy 25, 451–457 (2023).

14. Li, H., Song, W., Li, Z. & Zhang, M. Preclinical and clinical studies of CAR-NK-cell therapies for malignancies. Front. Immunol. 13, (2022).

15. Vivier, E., Ugolini, S., Blaise, D., Chabannon, C. & Brossay, L. Targeting natural killer cells and natural killer T cells in cancer. Nat Rev Immunol 12, 239–252 (2012).

16. Gong, Y. et al. NKG2A genetic deletion promotes human primary NK cell anti-tumor responses better than an anti-NKG2A monoclonal antibody. Molecular Therapy 32, 2711– 2727 (2024).

17. André, P. et al. Anti-NKG2A mAb Is a Checkpoint Inhibitor that Promotes Anti-tumor Immunity by Unleashing Both T and NK Cells. Cell 175, 1731–1743.e13 (2018).

18. Rölle, A., Jäger, D. & Momburg, F. HLA-E Peptide Repertoire and Dimorphism— Centerpieces in the Adaptive NK Cell Puzzle? Front. Immunol. 9, (2018).

19. Lanier, L. L. DAP10- and DAP12-associated receptors in innate immunity. Immunol Rev 227, 150–160 (2009).

20. Correia, M. P. et al. Distinct human circulating NKp30+FcεRIγ+CD8+ T cell population exhibiting high natural killer-like antitumor potential. Proc Natl Acad Sci U S A 115, E5980–E5989 (2018).

21. Lanier, L. L., Yu, G. & Phillips, J. H. Analysis of Fc gamma RIII (CD16) membrane expression and association with CD3 zeta and Fc epsilon RI-gamma by site-directed mutation. The Journal of Immunology 146, 1571–1576 (1991).

22. Zhang, T., Scott, J. M., Hwang, I. & Kim, S. Cutting Edge: Antibody-Dependent Memory-like NK Cells Distinguished by FcRγ Deficiency. The Journal of Immunology 190, 1402–1406 (2013).

23. Schlums, H. et al. Cytomegalovirus infection drives adaptive epigenetic diversification of NK cells with altered signaling and effector function. Immunity 42, 443– 456 (2015).

24. Shemesh, A. et al. Diminished cell proliferation promotes natural killer cell adaptive-like phenotype by limiting FcεRIγ expression. Journal of Experimental Medicine 219, e20220551 (2022).

25. Liu, W. et al. FcRγ Gene Editing Reprograms Conventional NK Cells to Display Key Features of Adaptive Human NK Cells. iScience 23, 101709 (2020).

26. Sugawara, S. et al. Knockdowns of CD3zeta Chain in Primary NK Cells Illustrate Modulation of Antibody-Dependent Cellular Cytotoxicity Against Human Immunodeficiency Virus-1. AIDS Research and Human Retroviruses 40, 631–636 (2024).

27. Aguilar, O. A., Fong, L.-K., Ishiyama, K., DeGrado, W. F. & Lanier, L. L. The CD3ζ adaptor structure determines functional differences between human and mouse CD16 Fc receptor signaling. Journal of Experimental Medicine 219, e20220022 (2022).

28. Otano, I. et al. CD137 (4-1BB) costimulation of CD8+ T cells is more potent when provided in cis than in trans with respect to CD3-TCR stimulation. Nat Commun 12, 7296 (2021).

29. Sheppard, S. et al. The Murine Natural Cytotoxic Receptor NKp46/NCR1 Controls TRAIL Protein Expression in NK Cells and ILC1s. Cell Reports 22, 3385–3392 (2018).

30. Zhuang, X., Veltri, D. P. & Long, E. O. Genome-Wide CRISPR Screen Reveals Cancer Cell Resistance to NK Cells Induced by NK-Derived IFN-γ. Front Immunol 10, 2879 (2019).

31. Sheffer, M. et al. Genome-scale screens identify factors regulating tumor cell responses to natural killer cells. Nat Genet 53, 1196–1206 (2021).

32. Pech, M. F. et al. Systematic identification of cancer cell vulnerabilities to natural killer cell-mediated immune surveillance. eLife 8, e47362 (2019).

33. Shemesh, A., Pickering, H., Roybal, K. T. & Lanier, L. L. Differential IL-12 signaling induces human natural killer cell activating receptor-mediated ligand-specific expansion. Journal of Experimental Medicine 219, e20212434 (2022).

34. Aguilar, O. A. et al. MICB Genomic Variant Is Associated with NKG2D-mediated Acute Lung Injury and Death. Am J Respir Crit Care Med 209, 70–82 (2024).

35. van Hall, T. et al. Monalizumab: inhibiting the novel immune checkpoint NKG2A. J Immunother Cancer 7, 263 (2019).

36. Heatley, S. L. et al. Polymorphism in Human Cytomegalovirus UL40 Impacts on Recognition of Human Leukocyte Antigen-E (HLA-E) by Natural Killer Cells *. Journal of Biological Chemistry 288, 8679–8690 (2013).

37. Battin, C. et al. NKG2A-checkpoint inhibition and its blockade critically depends on peptides presented by its ligand HLA-E. Immunology 166, 507–521 (2022).

38. Memmer, S. et al. The Stalk Domain of NKp30 Contributes to Ligand Binding and Signaling of a Preassembled NKp30-CD3ζ Complex*. Journal of Biological Chemistry 291, 25427–25438 (2016).

39. Luo, C. et al. Efficacy and safety of new anti-CD20 monoclonal antibodies versus rituximab for induction therapy of CD20+ B-cell non-Hodgkin lymphomas: a systematic review and meta-analysis. Sci Rep 11, 3255 (2021).

40. Swain, S. M., Shastry, M. & Hamilton, E. Targeting HER2-positive breast cancer: advances and future directions. Nat Rev Drug Discov 22, 101–126 (2023).

41. Cheng, Y. et al. The anti-tumor effects of cetuximab in combination with VTX-2337 are T cell dependent. Sci Rep 11, 1535 (2021).

42. Nieto, Y. et al. Allogeneic NK cells with a bispecific innate cell engager in refractory relapsed lymphoma: a phase 1 trial. Nat Med 1–7 (2025) doi:10.1038/s41591-025-03640-8.

43. Gauthier, L. et al. Multifunctional Natural Killer Cell Engagers Targeting NKp46 Trigger Protective Tumor Immunity. Cell 177, 1701–1713.e16 (2019).

44. Kaulfuss, M. et al. The NK cell checkpoint NKG2A maintains expansion capacity of human NK cells. Sci Rep 13, 10555 (2023).

45. Marçais, A. et al. High mTOR activity is a hallmark of reactive natural killer cells and amplifies early signaling through activating receptors. eLife 6, e26423.

46. Rölle, A., Meyer, M., Calderazzo, S., Jäger, D. & Momburg, F. Distinct HLA-E Peptide Complexes Modify Antibody-Driven Effector Functions of Adaptive NK Cells. Cell Reports 24, 1967–1976.e4 (2018).

47. Villard, M. et al. NK Cell Immaturity and NKp30 Expression Positively Correlate with Clinical Outcome in Multiple Myeloma Patients from the IFM2009 Clinical Trial. European Journal of Immunology 55, e202451191 (2025).

48. Chretien, A.-S. et al. NKp30 expression is a prognostic immune biomarker for stratification of patients with intermediate-risk acute myeloid leukemia. Oncotarget 8, 49548–49563 (2017).

